# αS oligomers generated from polyunsaturated fatty acid and dopamine metabolite differentially interact with Aβ to enhance neurotoxicity

**DOI:** 10.1101/2021.08.08.455587

**Authors:** Shailendra Dhakal, Jhinuk Saha, Courtney E. Wyant, Vijayaraghavan Rangachari

## Abstract

It is increasingly becoming clear that neurodegenerative diseases are not as discrete as originally thought to be but rather display significant overlap in histopathological and clinical presentations. For example, nearly half of the patients with Alzheimer disease (AD) and synucleinopathies such as Parkinson disease (PD) show symptoms and pathological features of one another. Yet, the molecular events and features that underlie such comorbidities in neurodegenerative diseases remain poorly understood. Here, inspired to uncover the molecular underpinnings of the overlap between AD and PD, we investigated the interactions between amyloid-β (Aβ) and α-synuclein (αS), aggregates of which form the major components of amyloid plaques and Lewy bodies, respectively. Specifically, we focused on αS oligomers generated from the dopamine metabolite called dihydroxyphenylacetaldehyde (DOPAL), and a polyunsaturated fatty acid docosahexaenoic acid (DHA). Both αS oligomers showed structural and conformational differences confirmed by their disparity in size, secondary structure, susceptibility to proteinase K digestion and cytotoxicity. More importantly, the two oligomers differentially modulated Aβ aggregation. While both oligomers inhibited Aβ aggregation to varying extents, they induced structurally different Aβ assemblies. Furthermore, Aβ seeded with DHA-derived αS oligomers showed greater toxicity than DOPAL-derived αS oligomers in SH-SY5Y neuroblastoma cells. These results provide insights into the interactions between two amyloid proteins with empirically distinctive biophysical and cellular manifestations, enunciating a basis for potentially ubiquitous cross-amyloid interactions across many neurodegenerative diseases.

## Introduction

It is increasingly becoming clear that neurodegenerative diseases defined by amyloid depositions are not discrete but show significant overlaps both clinically and histopathologically [1–3]. This is particularly the case with sporadic or familial Alzheimer disease (AD) and Parkinson disease (PD), where the up to 50% of patients from one pathology show the symptoms of the other [4]. In familial AD associated with the mutations in *APP, presenilins-1* and *2* genes, deposits of amyloid-β (Aβ) are often observed alongside Lewy bodies that involve the amyloid deposits of the protein α-synuclein (αS) [5, 6]. PD along with multiple system atrophy (MSA), and dementia with Lewy bodies (DLB) etc., involve the amyloid deposits of αS collectively called, synucleinopathies [7][8, 9]. Despite the pathological overlaps and co-morbidities, the molecular reasons behind such co-pathological presentations in neurodegenerative diseases remain unclear. Deciphering the molecular underpinnings of co-pathologies will require a deeper understanding of the cross-talks between multiple amyloidogenic proteins.

Aβ (40 and 42) is primarily generated in the extracellular space upon sequential cleavage of amyloid precursor protein (APP) by membrane bound aspartyl proteases, β-secretase and γ-secretase. Though Aβ42 is more aggressive than Aβ40 in terms of aggregation and toxicity, both forms of the protein are present in the plaque deposits. Implicated primarily in AD, Aβ aggregates are also observed in many other neurodegenerative pathologies including PD, amyotrophic lateral sclerosis (ALS), frontotemporal lobar degeneration (FTLD), and prion disease [10–14]. Similarly, αS involvement is also not limited to PD but also observed in a wide spectrum of synucleinopathies and in AD and ALS [15–18]. Previous clinical studies have shown the co-occurrence of αS and Aβ aggregates within the neocortical regions during the late stages of neurodegenerative pathologies. Brain autopsies has also revealed overlapping DLB/PD cases in clinically confirmed AD cases with characteristic Aβ/tau aggregates [19]. These observations have propelled *in vitro* studies, which show direct interactions between monomeric αS and Aβ resulting in the cross-seeding effects leading to fibrils formation [20]. Similarly, our lab has also shown the cross interaction of dopamine-derived αS oligomers (DSOs) with Aβ42 monomers resulting in higher molecular weight oligomers formation [21]. A more recent study has shown that αS oligomers generated from dopamine and those derived from the polyunsaturated fatty acid, docosahexaenoic acid (DHA) show mesoscale differences [22]. In addition, the two oligomers exhibited differential cross interactions with tau leading to diverse biophysical characteristics and cytotoxic events [22]. The variations in oligomer conformations and fibril polymorphisms among amyloid species is emerging to be a key contributor of phenotypes and clinical variations [23–25], making the investigations involving a diverse set of oligomers imperative.

Based on the cues from the effect on tau, we investigated the effects of 3,4-dihydroxyphenylacetaldehyde (DOPAL) and DHA-derived αS oligomers (from here on, DOPAL-SOs and DHA-SOs, respectively) on Aβ aggregation. We hypothesized that these oligomers of αS may cross-interact with Aβ monomers causing alteration of the latter aggregation and toxicity. DOPAL is a monoamine oxidase (MAO) derived metabolite of dopamine which is toxic to the neurons at physiological concentration [22, 26–28]. It is shown to be associated with damage of synaptic vesicles. Moreover, injection of DOPAL in rats led to loss of dopaminergic neurons with accumulation of αS oligomers [29]. The generation of αS oligomers in presence of DOPAL involves Schiff’s base and Michael addition with the free amines of lysines in αS to form covalent adducts [28] (**Fig 1a**). On the other hand, DHA is a polyunsaturated fatty acid with diverse physiological functions and is a major component of myelin sheath. Upon incubation with αS monomers, DHA is shown to form both covalent adducts and engage in non-covalent interactions to generate oligomers with α-helical characteristics [30] (**Fig 1b**). However, the structural transition to random coil occurs with the decrease in DHA content in the oligomers [30, 31]. In this study, we show that DOPAL-SOs and DHA-SOs are conformationally distinct with different biophysical characteristics. Our results indicate that the two αS oligomers differentially interact with Aβ to inhibit its aggregation and generate species with distinct secondary characters, biophysical properties. More importantly, the two oligomers promote Aβ aggregate species that enhances cytotoxicity than the unseeded Aβ control demonstrating the ability of the oligomers to induce distinctive cellular responses.

**Figure 1.**
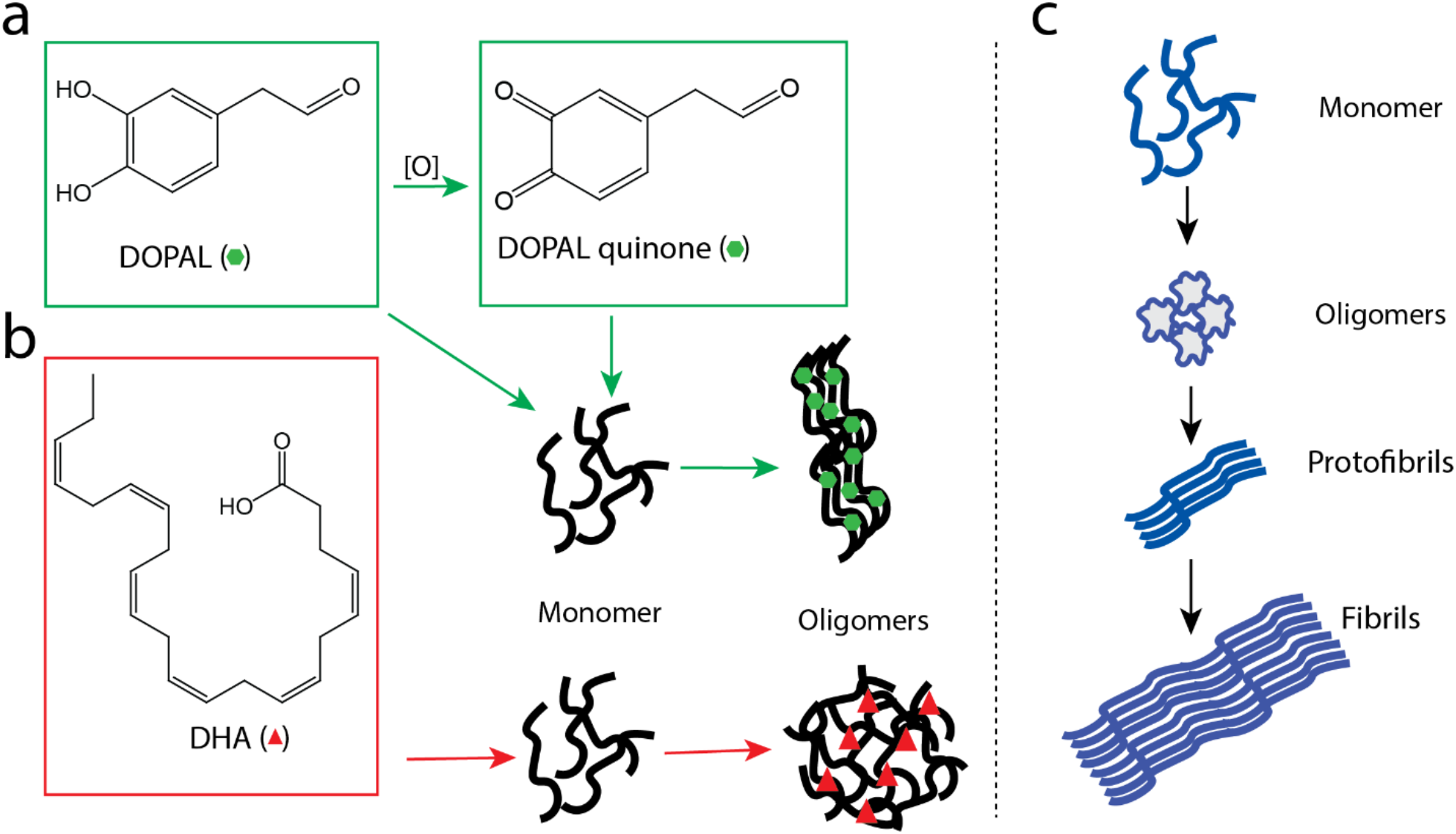
Schematic diagram showing on-pathway and off-pathway αS aggregation. a) DOPAL forms DOPAL quinone in the oxidizing environment which interacts with αS monomers both covalently and non-covalently resulting in oligomers formation. In the reduced form, it can form Schiff’s base with αS monomers to form oligomers. b) DHA can interact with αS monomers through oxidative modification at four methionine residues and covalent carbonyl adduct formation at the protein side chains to form structurally distinct αS oligomers. c) Schematics of on-pathway αS aggregation resulting in oligomers, protofibrils and fibrils formation.

## Results

### DOPAL and DHA-derived αS oligomers show biophysical differences

As mentioned above, DOPAL and DHA are known to form covalent adducts with αS generating oligomers which we suspected to be conformationally discrete to one another with distinctive properties. The chemistry of DOPAL is fairly well understood; in the reduced form, DOPAL can form a Schiff’s base with free amines in αS and upon oxidation, it is converted into its quinone form, which covalently or non-covalently interact with αS monomers resulting in oligomers formation [28]. Similarly, DHA is known to covalently interact with αS monomers and promote oligomerization [22, 31]. These are characterized as “off-pathway” oligomers of αS that are not only temporally different from “on-pathway” counterparts but also structurally [31,32] **(Fig 1)**. To investigate the properties of DOPAL-SOs and DHA-SOs, seed-free αS monomers were incubated with DOPAL and DHA in separate reactions with orbitally shaking at physiological temperature.

After 24h, DOPAL-SOs were fractionated using size exclusion chromatography (SEC) which showed two distinguishable peaks between fractions 16-20 and 21-23 **(Fig 2a)**. The first peak corresponding to fractions 17 and 18, which are the major fractions of the first peak showed the presence of mixture of oligomers ranging 50-260 and 36-260 kDa disperse bands, respectively (***inset*, Fig 2a**). The second peak in the fraction 21-23 revealed the presence of major monomeric and dimeric species in SDS-PAGE gel (data not shown). Size analysis of the fractions 17 and 18 by dynamic light scattering (DLS) showed monodispersed peaks with a mean hydrodynamic diameter ranging from 10-40 nm, while the corresponding monomers showed 3-6 nm hydrodynamic diameter indicating presence of some low molecular weight oligomeric species possibly dimers or trimers (**Fig 2b**). This is also captured in the autocorrelation function with the correlation coefficient for fraction 17 showing a slightly longer correlation times as compared to fraction 18 reflecting a slightly larger size of the former than the latter (**Fig 2c**). SEC fractionation of DHA-SOs also showed a large peak near the excluded volume ranging from fractions 15 to 20 along with a small peak near the fraction numbers 25-30 corresponding to the monomers (**Fig 2d**). Immunoblotting of the fractions 16 and 17 revealed wide distribution of oligomeric species populated throughout the higher molecular weight region as compared to DOPAL-SOs (***inset*, Fig 2d**). Corresponding DLS also showed wide size distribution with the mean hydrodynamic diameter ranging from 20-80 nm **(Fig 2e)**. Among the two, fraction 16 showed slightly large size as expected as compared to fraction 17 (**Fig 2e and f)**.

**Figure 2.**
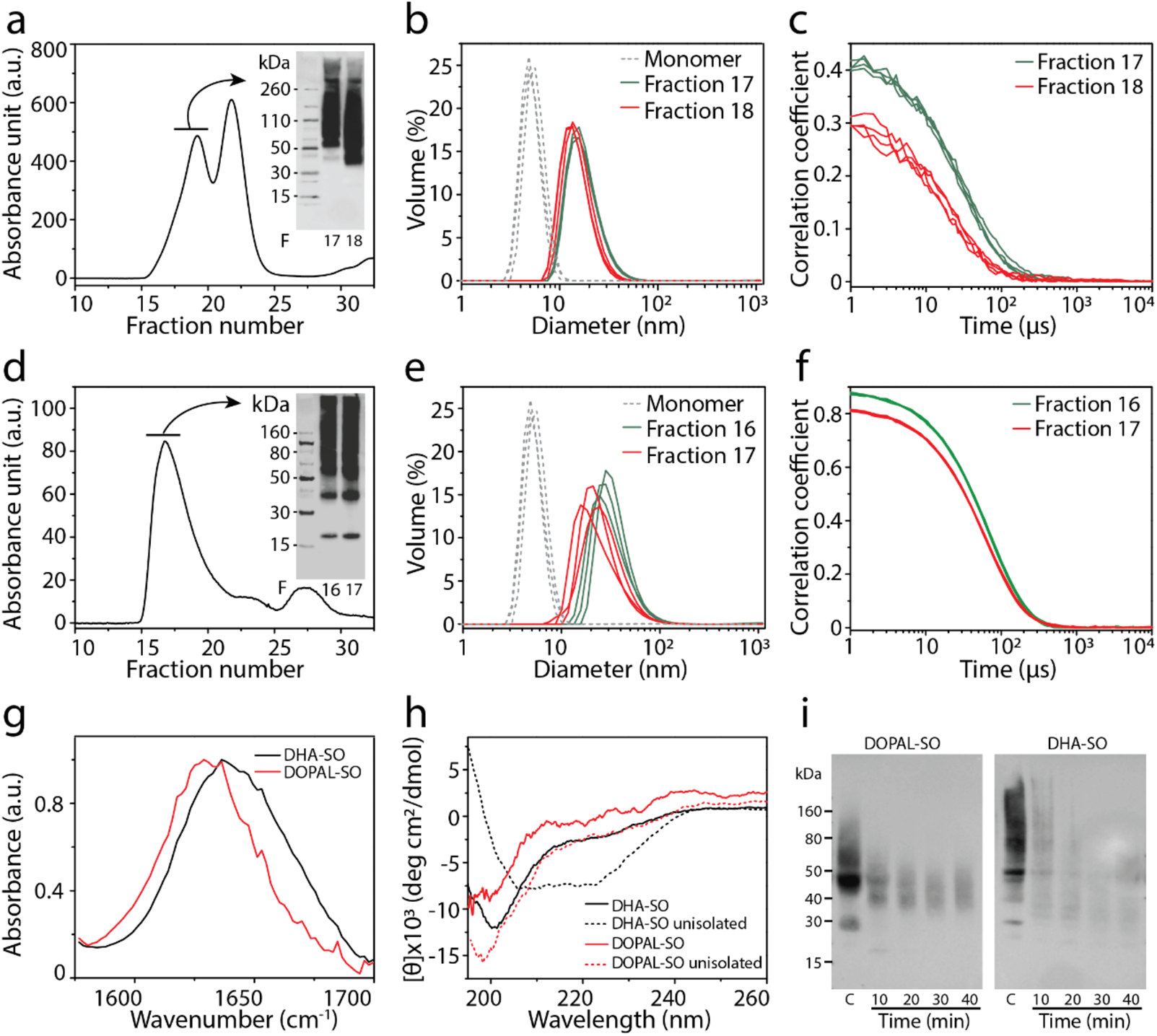
Isolation and characterization of DOPAL-SOs and DHA-SOs. a) Size exclusion chromatogram of DOPAL-SOs along with immunoblots (inset) of fractions 17 and 18 (indicated by arrow) using αS Syn211 antibody. b-c) Hydrodynamic diameter of monomers and respective oligomers using DLS (b) and correlation coefficient (derived from autocorrelation function) of the respective oligomers plotted as a function of time (c). d) Size exclusion chromatogram of DHA-SOs with immunoblots (inset) of fractions, 16 and 17. e-f) Hydrodynamic diameter of monomers and oligomeric fractions represented as volume percentage (e) and their correlation coefficient as a function of time (f). g-h) Normalized FTIR and far-UV CD spectra of isolated DOPAL-SOs and DHA-SOs. h) black dotted lines and red dotted lines represent DHA-SOs and DOPAL-SOs respectively prior to SEC isolation. i) Immunoblot showing PK digestion data of DOPAL-SOs and DHA-SOs at 10, 20,30 and 40 minutes of incubation; C represents control oligomers without PK treatment.

Secondary structure of these oligomers was studied using Fourier-transform infrared spectroscopy (FTIR) and far-UV circular dichroism (CD) spectroscopy. FTIR spectra revealed the random coiled structure of DOPAL-derived αS oligomers with peak at ~1640 nm, while DHA generated αS oligomers showed peak at 1640 nm and 1650 nm indicating presence of random coiled structure with slightly α-helical characteristics **(Fig. 2g)**. CD spectra of the oligomers showed slightly different secondary structures. Both SEC-isolated **(*red*)** and non-isolated **(*red dash lines*)** DOPAL-SOs exhibited corresponding minima at 195 nm characteristics of random coiled structure **(Fig. 2h)**, which is similar to the dopamine-derived oligomers [21]. DHA-derived oligomers showed different secondary structure based on the degree of DHA associated with the protein. While SEC-isolated oligomers without free DHA showed mainly random coiled structure with a minimum at 195 nm (***black*; Fig 2h**), when complexed with DHA prior to SEC isolation, the sample displayed α-helical characteristics with a minimum at 208 and a shoulder at 222 nm **(*black dash lines*; Fig 2h**), which is the form widely reported in the literature [22, 30, 31, 33–35]. Finally, enzymatic stability of αS oligomers were probed by proteinase K (PK) digestion. Stability of the oligomers were assessed by disappearance of oligomeric band in immunoblots upon treatment with PK, digested at specific time intervals of 10, 20, 30 and 40 mins. The samples were then run on SDS PAGE gel and visualized by immunoblot using monoclonal anti-αS (Syn211) antibodies. Upon doing so, the intensities of the bands for both DOPAL and DHA-derived αS oligomers were diminished significantly compared to the controls within 10 minutes of PK digestion (**Fig 2i**). However, the intensity of DHA-derived αS oligomers is found be much lower than the intensity of DOPAL-derived αS oligomers (**Fig 2i**). This indicates higher susceptibility of DHA oligomers for PK digestion as compared to DOPAL-SOs **(Fig. 2i)**. The higher stability of DOPAL-SOs could result from the covalent Schiff’s base adducts present within the oligomers which may provide tighter binding interactions leading to a more compact structure resistant to enzymatic cleavage. On the contrary, DHA oligomers, which are formed by both covalent and non-covalent interactions seem to less stable. Nevertheless, the data clearly point out that DOPAL-SOs and DHA-SOs are structurally and conformationally different from one another.

### Both DOPAL and DHA-derived αS oligomer differentially modulate Aβ aggregation

We further investigated the ability of DOPAL-SOs and DHA-SOs to cross-seed Aβ42. To do so, 0.75 μM of SEC fractionated DOPAL-SOs (fraction 17 or 18) were incubated with seed-free, 15 μM Aβ42 monomers and monitored for 48 hours by thioflavin-T (ThT) fluorescence. Aβ42 control in the absence of oligomer seeds showed sigmoidal ThT fluorescence curve with 10 hours of aggregation lag time (**Fig 3a**). However, the addition of either fraction 17 or 18 of DOPAL-SOs decreased the aggregation by increasing the lag time of Aβ aggregation to ~35 hours along with decrease in plateau intensity of ThT fluorescence **(Fig. 3a)**. Aliquots of the samples from all three reactions were removed at given time points and were analyzed by immunoblotting using Ab5 monoclonal antibody for Aβ42. Up to 20 h, the samples incubated with DOPAL-SOs predominantly showed the presence of monomers in the immunoblots while the control Aβ showed high molecular weight species after 20 h (**Fig 3b**).

**Figure 3.**
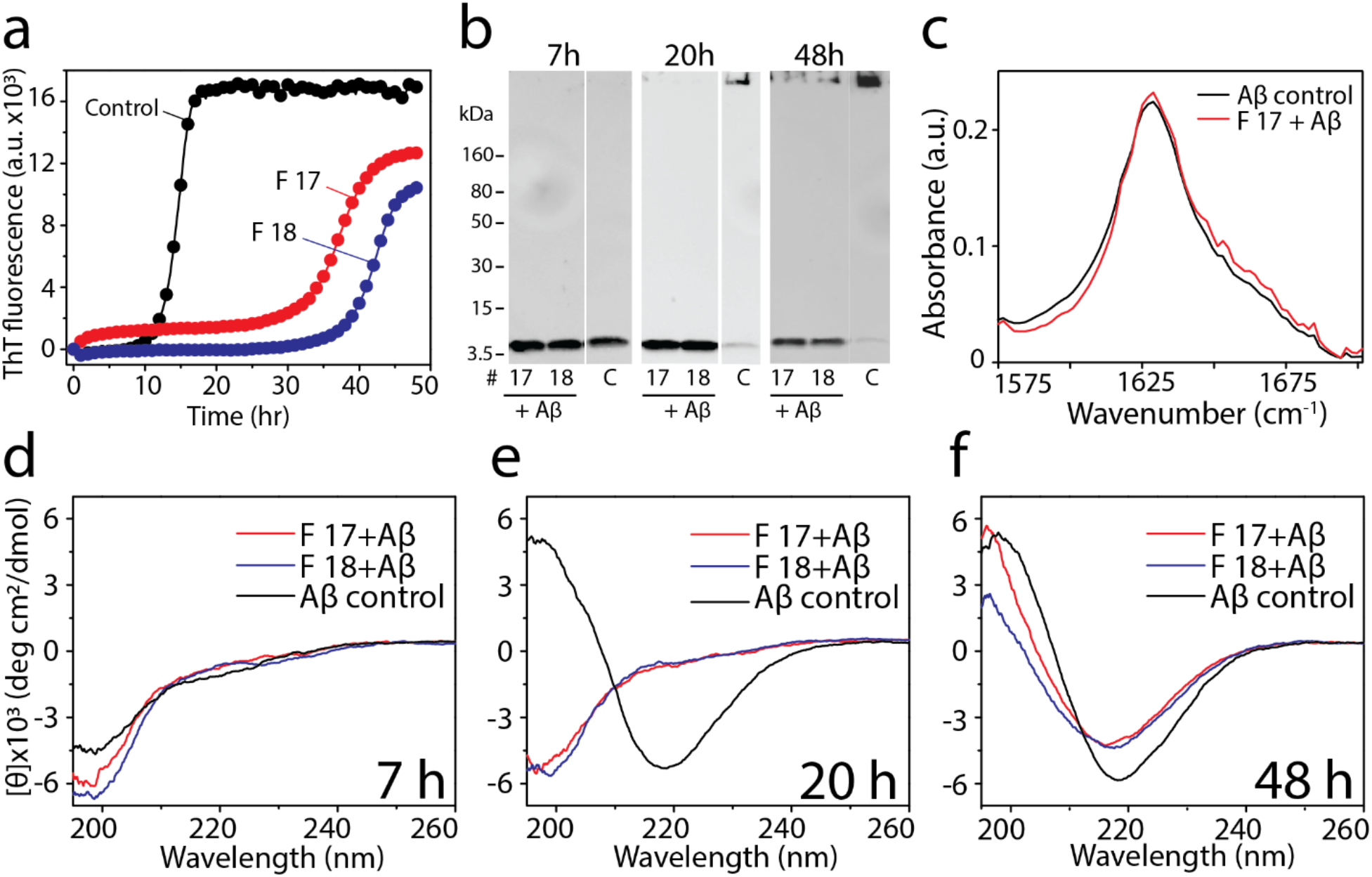
Interaction of DOPAL-SOs with Aβ. a) ThT fluorescence kinetics of 15 μM Aβ monomers (●) alone and in the presence of 0.75 μM DOPAL-SO corresponding to 17 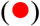 and 18 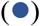 fraction number in 20 mM tris buffer pH 8.0. b) Immunoblot of ThT reactions (figure 3,a) at 7 h, 20 hours and after 48 h, C indicates Aβ control sample. c) FTIR analysis of reactions containing Aβ monomer (15 μM) alone and in presence of 0.75 μM DOPAL-SOs (fraction 17) after 48 h. d-e) CD spectra of ThT reactions (figure 3,a) at 7 h, 20 h and after 48 h respectively.

After 48 h, the control Aβ aggregated to form fibrils that failed to enter the gel but the samples with DOPAL-SOs showed both monomers and some fibrils mirroring the ThT data (**Fig 3b**). In parallel, conformational changes of the samples were monitored by far-UV CD. The CD spectra for the Aβ control also showed a conformational change from a random coil at 7 h of incubation to a β-sheet after 20h with characteristic minima at 220 nm and maxima at 198 nm (***black*; Fig 3c-e**). Co-incubation of fractions 17 and 18 of isolated DOPAL-SOs with Aβ remained in a random coil conformation up to 20 h before converting to a β-sheet after 48 h (***red and blue*; Fig 3c-e**). The fibrils formed at the end of the reaction after 48 h of incubation was sedimented and analyzed with FTIR which showed parallel β-sheet rich structure for both the co-incubated reactions and control with peak centered at 1630 cm^-1^ (**Fig 3c**). These results indicate that DOPAL-SOs tend to delay the aggregation of Aβ42 but form a similar β-sheet secondary structure of fibrils as the control **(Fig 3c)**. However, these observations do not exclude the possibility of having dissimilar molecular arrangements of the structure as CD and FTIR are inconspicuous to such atomic level changes.

Next, we investigated the interactions between DHA-SOs and Aβ42 by incubating 0.75 μM each of DHA-SOs fraction 16 and 17 with freshly prepared, seed-free 15 μM Aβ42 monomers at physiological temperature. ThT fluorescence indicated that the Aβ aggregation was attenuated by the oligomers based on the increase in the lag times observed **(Fig. 4a)**. This was also evident from the immunoblots which showed inhibition of aggregation by both fractions of DHA-SOs as opposed to the control Aβ in the absence of oligomers **(Fig 4b)**. Fibrillar species were not detected in the reactions containing Aβ monomers co-incubated with DHA-SOs. In far-UV CD acquired in parallel, conformational changes of Aβ in the presence of DHA-SOs with monomer showed a faster conversion from random coiled to β-sheet structure than the control without the oligomers **(Fig 4d-f)**. However, the DHA-SO co-incubated samples showed spectra that were deviated from ideal β-sheet with minimum at 220 nm but contained a partial α-or 3_10_-helical characteristics with a negative shoulder at 208 nm (***red and blue*; Fig 4d-f**) unlike those with DOPAL-SOs. FTIR analysis of the samples after 48 hours confirmed the presence of β-sheet structure for the Aβ control, while combination of α-helical and β-sheet characteristics for the reaction similar to the CD **(Fig 4c)**. Together, these data suggest that DHA-SOs also inhibit Aβ aggregation but do so by converting Aβ to a conformation containing a mixture of α-helical and β-sheet secondary structures.

**Figure 4.**
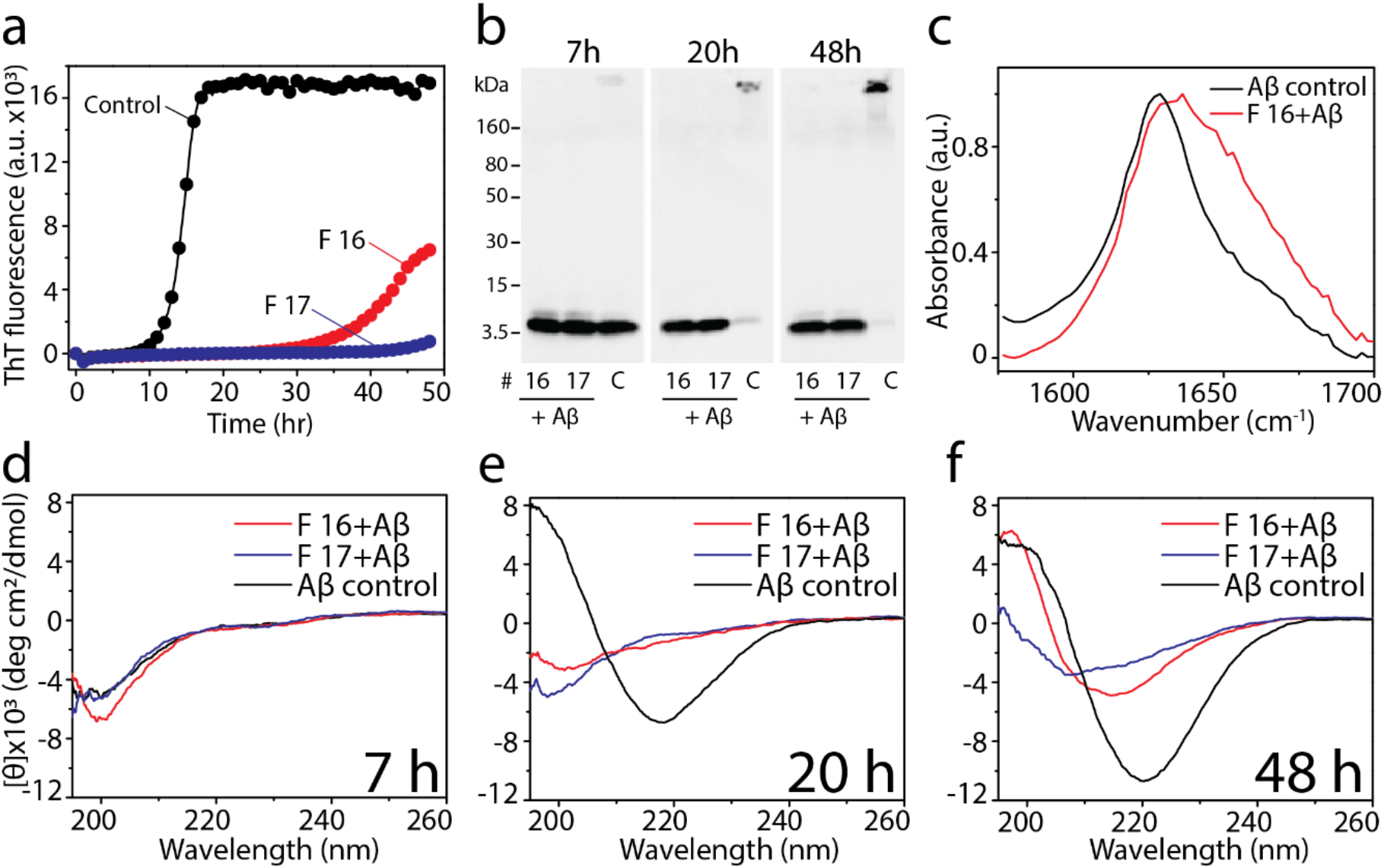
Interaction of DHA-SOs in presence of Aβ monomers. a) ThT fluorescence of 15 μM Aβ monomer alone (●) and in the presence of 0.75 μM DHA-SOs corresponding to fraction number 16 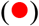 and 17 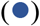. b) Immunoblotting of ThT reactions (Figure 4a) at 7 h, 20 h and after 48 h using Ab5 monoclonal antibody, C represents the Aβ control reaction. c) FTIR analysis of 15 μM Aβ monomer alone and in the presence of 0.75 μM DHA-SOs after 48 h. d-e) CD spectra of ThT reactions (figure 4a) at 7 h, 20 h and 48 h.

### Aβ aggregates generated from seeding DOPAL and DHA-derived αS oligomers show enhanced neurotoxicity

It is clear from the data presented thus far that both DOPAL-SOs and DHA-SOs have subtle differences that seem to manifest in the way they seed Aβ aggregation. To see whether these differences manifest in their cellular toxicity, their effects were investigated for cell viability in neuroblastoma cells. Firstly, the reaction containing 15 μM Aβ was incubated with and without 0.75 μM DOPAL-SOs or DHA-SOs at 37 °C for 48 hours. Aliquots of reaction were diluted two-folds in DMEM/F-12 and control samples containing 7.5 μM Aβ monomer, and 325 nM DOPAL-SOs or DHA-SOs were added to SH-SY5Y neuroblastoma cells seeded 24 hours prior to experiment. The toxicity of these species in neuroblastoma cells was examined by using XTT cytotoxicity assay (described in Methods). The results indicate the greater toxicity of Aβ species formed in the presence of the oligomers as compared to their respective controls (**Fig 5**). Aβ monomers showed least toxicity among all species, while DHA-SOs in presence of Aβ showed highest toxicity (**Fig 5**). Together, the data suggest that the cross-seeding of αS oligomers with Aβ augments toxicity.

**Figure 5.**
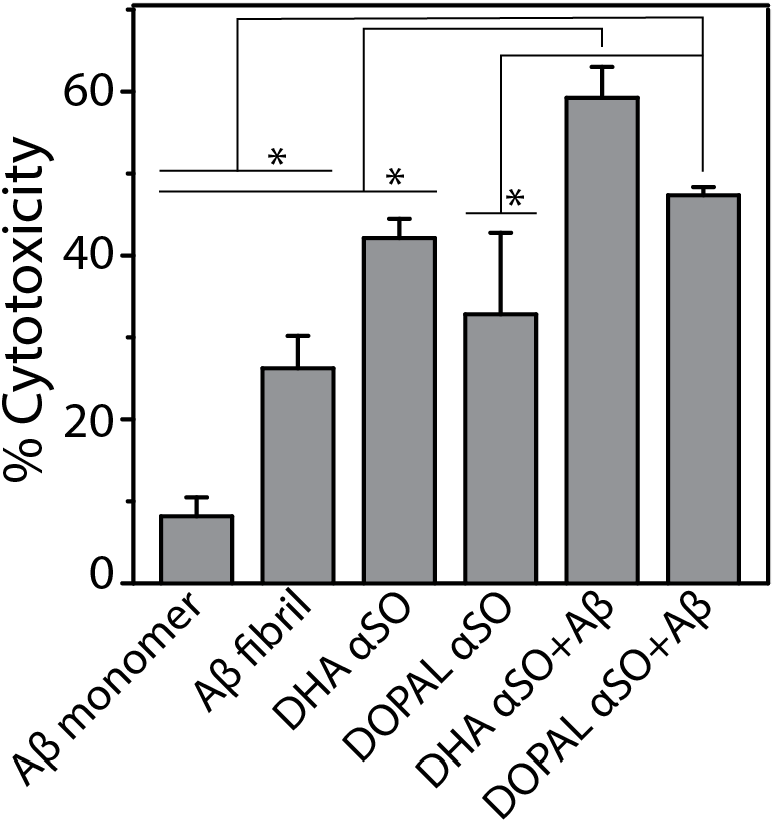
Cytotoxicity of DOPAL-SOs and DHA-SOs with and without Aβ using XTT in mammalian SH-SY5Y neuroblastoma cells. Data were obtained in triplicates, * represents p<0.05 using one-way ANOVA analysis.

## Discussion and conclusions

In this study, we generated two physiologically relevant and conformationally distinct αS oligomers, DOPAL-SOs or DHA-SOs, which showed subtle biophysical differences and chemically different modes of interaction with Aβ. The results show that the two oligomers interact with and delay the rate of Aβ aggregation to different extents. More importantly, the modulation of Aβ aggregation by the two αS oligomers was accompanied by the enhancement of cytotoxicities as compared to either Aβ alone or DOPAL-SOs or DHA-SOs. Direct interaction of monomeric αS and Aβ, oligomeric αS and tau in relevance to AD and synucleinopathies has been shown in the previous studies [20, 22]. Despite the preponderance of AD in patients with synucleinopathies, the interplay between specific αS oligomers and Aβ, key constituents of Lewy bodies and plaques respectively, has remained unexplored. The conformational diversity among amyloid oligomers and their cellular mechanisms have been well established, but not in the context of cross-interactions with other amyloid proteins. In this study, we sought to understand the dynamics and cellular effects of cross-interactions between and αS oligomers and Aβ that has remained elusive. We have previously generated and characterized dopamine-derived αS oligomers (DSOs) capable of homotypic and heterotypic interactions [21]. In this study, we focused on the more reactive and toxic intermediate of dopamine metabolism, DOPAL. Dopamine is oxidized in the cytoplasm by MAO to toxic DOPAL, which is shown to be upregulated in PD [32]. We also used DHA fatty acid, a major component of myelin which covalently binds to αS and generate oligomers [30, 36].

We observed that DOPAL-SOs and DHA-SOs show subtle yet important differences; DHA-SOs have a larger hydrodynamic radius than DOPAL-SOs, both have distinct secondary structures and show differences in stability towards proteinase K digestion. These differences are clearly manifested in the way they interact with Aβ monomers. While both oligomers delayed Aβ aggregation, DHA-SOs showed more effective inhibition than DOPAL-SOs. The inhibition of Aβ aggregation kinetics by DOPAL-SOs still resulted in the formation of aggregates with β-sheet characteristics, however, DHA-SOs attenuated Aβ aggregation resulting in the formation of Aβ species with mixture of random coiled, α-helical and β-sheet secondary structure. More importantly, these oligomers were toxic to the neuroblastoma SH-SY5Y cells by themselves, but their interaction with Aβ led to greater degree of toxicity possibly due to formation of polymorphic Aβ species [37]. Among all the samples, DHA-SO seeded Aβ displayed the highest degree of toxicity which may be due to enhanced surface hydrophobicity of the oligomers and resulting species formed after interaction as reported [38, 39].

In sum, the study presented here highlights the molecular basis of interplay and synergism between Aβ and αS oligomers that has not been known before. These results help unravel the principle behind cross-interactions among amyloid proteins is that depending on the conformation and properties of oligomers of αS, Aβ monomers seeded by these oligomers may generate a polymorph of Aβ fibrils with distinctive biophysical and cellular properties as schematically depicted in **Fig 6**. These mechanistic insights also help deepen our understanding of the clinical and pathological overlaps observed in AD and synucleinopathies, the one that could arise from the interaction between Aβ and αS. Further details of the mechanisms will continue to emerge that could precisely decipher the dynamic interplay between monomers, oligomers, and fibrils of Aβ and αS.

**Figure 6.**
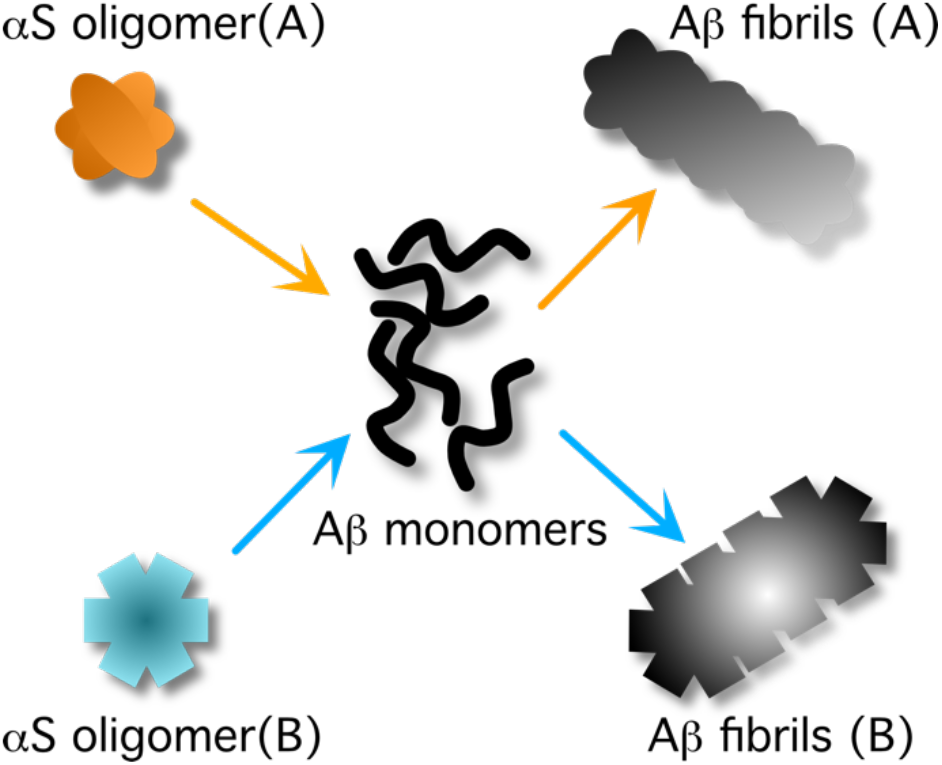
Generalized schematic summary from the results obtained.

## Materials and methods

### Materials

DOPAL and DHA were purchased from Cayman Chemicals and Sigma-Aldrich respectively. Monoclonal antibody, Syn 211 (αS specific) was purchased from Millipore Sigma, while Ab5 was a generous gift from Dr. Levites (University of Florida). All other routine laboratory chemicals and consumables were purchased from Thermo scientific.

### Recombinant protein expression and purification

Recombinant expression and purification of αS and Aβ were carried out following previously established protocol [40, 41]. Briefly, αS and Aβ42 plasmid were transformed and grown in *E. coli* BL21 (DE3) cells, and protein expression was induced with 1 mM IPTG. For αS, cells were harvested and lysed in lysis buffer (20 mM Tris, 50 mM NaCl, 10 mM imidazole, pH 8.0) in presence of PMSF and sonicated using Misonix XL-2000 for eight cycles (45 sec brust with 1 min rest). Lysate was centrifuged at 15,000 x g and supernatant was loaded into Ni-NTA affinity column. Protein was washed using two sets of wash buffers-wash 1 containing 20 mM Tris, 50 mM NaCl, 20 mM imidazole and wash 2 containing 20 mM Tris, 50 mM NaCl, 75 mM imidazole at pH 8.0. The column was further subjected to elution buffer containing 20 mM Tris, 50 mM NaCl, 250 mM imidazole at pH 8.0. The eluted protein was dialyzed against distilled water for 2 cycles each of 2 hours and lyophilized. The lyophilized protein was stored at −80 °C. Protein was resuspended 50 mM NaOH in Millipore water, incubated for 30 minutes and subjected to size exclusion chromatography (SEC) in 20 mM Tris pH 8.0 to obtain monomeric αS. For Aβ, harvested cells were lysed in lysis buffer (20 mM Tris Buffer, 1mM EDTA, pH 8.00) and sonicated for 10 cycles (30 sec brust with 1 min rest) followed by centrifugation at 10000 x g for 10 min. Pellets were collected and 2 more rounds of sonication cycle was carried out with 8 cycles and 6 cycles respectively followed by centrifugation at 15000 x g for 10 min between each step. The pellet was then dissolved in 4 M urea and sonicated for another 6 cycles. The lysate in urea was centrifuge at 15000 x g for 10 minutes. Supernatant was collected and passed through a 0.2 µm to remove any debris. The filtrate was subjected to HPLC and the obtained Aβ was lyophilized for long term storage. To freshly purify Aβ monomers, lyophilized Aβ was incubated in 10mM NaOH and subjected to SEC.

### Preparation of DOPAL and DHA-derived αS oligomers

DOPAL-SOs were prepared by incubating 50 μM monomeric αS with 1 mM DOPAL in 20 mM Tris buffer pH 8.0. Similarly, DHA-SOs were prepared by incubating 50 μM monomeric αS with 2.5 mM DHA in 1X PBS. Both reactions were incubated at 37 °C in an incubator shaker at 700 rpm for 24 hours and subjected to SEC in 20 mM Tris buffer pH 8.0. Isolated oligomers concentration was calculated and characterized using DLS.

### SDS-PAGE and western blotting

Sample aliquots of isolated αS oligomers and their cross-seeding reaction with Aβ were subjected to SDS-PAGE and immunoblotting using monoclonal anti-αS antibody, clone syn211 (Millipore sigma), monoclonal Ab5 antibody as described previously [42]. Briefly, all the samples were mixed with 4x Laemmli sample loading buffer and loaded onto SDS-PAGE Biorad Mini-PROTEAN® 4-20 % precast gel. For immunoblotting, gels were transferred on to a 0.45 μm Amersham Protran Premium nitrocellulose membrane (GE life sciences) and blot was boiled for one minute in 1X PBS. Blot was incubated overnight in the blocking buffer (5% non-fat dry milk, 1% Tween ®-20 in 1X PBS) followed by primary antibodies against αS or Aβ, and anti-mouse secondary antibodies each for two hours. Blot images were acquired on GelDoc molecular imager (Bio-Rad) after treating with ECL reagent.

### Dynamic light scattering

Dynamic light scattering (DLS) experiments were carried out in a Zetasizer Nano S instrument (Malvern, Inc.). The data were acquired with 70 μL sample volume by averaging 3 runs each of 10 s with pre-equilibration time of 30 s. All the parameters were determined, autocorrelation function was plotted as a function of time, diameter was calculated using volume (%) function, and plotted in origin 8.5.

### Circular dichroism

Circular dichroism (CD) spectra of respective samples were measured in 20 μM Tris buffer pH 8.0 in the far UV region in Jasco J-815 spectrophotometer (Jasco MD) using the established protocol [43].

### Fourier Transform Infrared (FTIR) spectroscopy

Samples of monomeric proteins, DOPAL and DHA-derived αS oligomers, and fibrils from the cross-seeding reaction were lyophilized and dissolved in D_2_O and measured in Cary 630 FTIR spectrometer. FTIR spectra were acquired with 1024 total scans from 1800 cm^-1^ to 1400 cm^-1^ at a resolution of 4 cm^-1^. Spectra were blank subtracted with D_2_O, normalized and plotted in origin 8.5.

### Thioflavin-T (ThT) binding assay

ThT aggregation kinetics was performed by incubating the samples with 10 μM ThT and data were monitored using BioTek Synergy H1 microplate reader at 37 °C. Samples excitation and emission was set as 452 nm and 485 nm respectively. Data were plotted as ThT fluoresncence versus time in origin 8.5.

### Proteinase K digestion

DOPAL-SOs and DHA-SOs (13.3 μg) were digested with 0.9 ng proteinase K (PK) diluted from a stock of 20 mg/mL (Ambion, Inc). Reactions were run at 37° C by shaking at 200 rpm and aliquots of the reaction were quenched with 0.5 mM PMSF at 10, 20, 30 and 40 minutes, respectively. Post quenching, 9.95 ng of reaction samples were run in a Invitrogen 4-12% SDS-PAGE gel and transferred onto an 0.2 µm nitrocellulose membrane. Protein bands were investigated by using anti-αS monoclonal antibody (Syn 211, Millipore Sigma) and anti-mouse horse radish peroxidase secondary antibody. The immunoblots were imaged with Super Signal West Pico Chemiluminescent Substrate kit (Thermo Fisher Scientific).

### Cell viability

Cell viability assay was carried out using 2,3-bis(2-methoxy-4-nitro-5-sulfophenyl)-5-[(phenylamino)carbonyl]-2H-tetrazolium hydroxide (XTT) in SH-SY5Y cells as described in our previously established protocols [40, 43]. Briefly, human neuroblastoma SH-SY5Y were maintained at 37 °C with 5.5 % CO_2_ in DMEM: F12 (1:1) media containing 10% FBS and 1% penicillin/streptomycin. Cells were plated in 96 well plates 24 hours prior to experiment and treated with respective samples. After incubating the samples for 24 hours prior, both the blank and experimental readings were acquired in BioTek Synergy H1 microplate reader.

## Acknowledgements

The authors would like to thank the following agencies for financial support: National Institute of Aging (1R56AG062292-01), National Institute of General Medical Sciences (R01GM120634), and the National Science Foundation (NSF CBET 1802793) to VR. The authors also thank the National Center for Research Resources (5P20RR01647-11) and the National Institute of General Medical Sciences (8 P20 GM103476-11) from the National Institutes of Health for funding through INBRE for the use of their core facilities.

## Author contributions

VR conceptualized the project; SD performed protein purification, biophysical experiments along with CEW, SD also conducted the cell culture experiments, and JS conducted proteinase K digestion and collected FTIR data. VR, SD and JS participated in intellectual discussions and manuscript writing and editing.

